# Molecular evolution in large steps - Codon substitutions under positive selection

**DOI:** 10.1101/510958

**Authors:** Qingjian Chen, Ziwen He, Ao Lan, Haijun Wen, Chung-I Wu

**Affiliations:** State Key Laboratory of Biocontrol, School of Life Sciences, Sun Yat-sen University, Guangzhou 510275, China; CAS Key Laboratory of Genomic and Precision Medicine, Beijing Institute of Genomics, Chinese Academy of Sciences, Beijing 100101, China; Department of Ecology and Evolution, University of Chicago, Chicago Illinois 60637, USA

## Abstract

Molecular evolution is believed to proceed in small steps. The step size can be defined by a distance reflecting physico-chemical disparities between amino acid (AA) pairs that can be exchanged by single 1 bp mutations. We show that AA substitution rates are strongly and negatively correlated with this distance but only when positive selection is relatively weak. We use the McDonald and Kreitman (MK) test to separate the influences of positive and negative selection. While negative selection is indeed stronger on AA substitutions generating larger changes in chemical properties of amino acids, positive selection operates by different rules. For 65 of the 75 possible pairs, positive selection is comparable in strength regardless of AA distance. However, the 10 pairs under the strongest positive selection all exhibit large leaps in chemical properties. Five of the 10 pairs are shared between hominoids and *Drosophila*, thus hinting at a common but modest biochemical basis of adaptation across taxa. The hypothesis that adaptive changes often take large functional steps will need to be extensively tested. If validated, molecular models will need to better integrate positive and negative selection in the search for adaptive signal.

## Introduction

It is generally accepted that natural selection favors incremental small-step changes. In the accompanying paper (<u>Chen and Wu</u>), we use physico-chemical distances between amino acids (AAs) as a measure of step size in evolution. When negative selection is the main driving force, similar AAs are indeed more likely to be exchanged. After all, a mutant must not be too different from the wild type in order to avoid elimination. This intuition is a key rule of neutral molecular evolution (<u>Kimura 1983</u>).

On the other hand, it is not obvious that positive selection should also favor small-step changes. The most common reference is Fisher’s geometric model (FGM) (<u>Fisher 1930</u>), whereby small-step changes would have a better chance of being advantageous than large-step ones. Nevertheless, a key element of FGM is still the avoidance of negative selection. In FGM, each mutation is assumed to be highly pleiotropic and large-step changes are likely to be deleterious for some phenotypes (<u>Wagner and Zhang 2011</u>).

An opposite argument for large-step evolution can be stated as follows: Natural selection, either positive or negative, can “discern” large-step changes better than small-step ones. For example, replacing isoleucine (Ile) with the chemically similar Valine (Val) would not alter the protein structure as much as its replacement by Arginine (Arg), which is very different chemically. In this view, an Ile ➔ Arg replacement may be either much worse, or much better, than the Ile ➔ Val replacement. Therefore, while negative selection would accept small-step changes (e.g., Ile ➔ Val), positive selection might in fact favor large-step ones (e.g., Ile ➔ Arg). The conventional wisdom is a postulate, not a fact.

Clearly, the arguments must be resolved by empirical means. The companion study (<u>Chen and Wu</u>) has shown that AA substitutions start to deviate from the small-step rule when negative selection becomes weaker and/or positive selection becomes stronger. In the same vein, this study aims to separate the two effects on the rate of AA substitutions.

## Results

### I. Amino acid distance in relation to the action of negative selection

We partitioned the conventional Ka (or Dn) measurement (number of non-synonymous substitutions per site) into 75 classes of substitutions, denoted Ki (i = 1, 75; (<u>Tang, et al. 2004</u>)). These 75 classes are AA substitutions that require only a 1 bp change. Ks (or Ds), the number of synonymous substitutions per site, is a separate class. It is reported that the rank order of Ki is highly correlated (R > 0.9) across taxa ranging from plants and invertebrates to mammals and primates (<u>Chen and Wu</u>). The main reason for this nearly universal correlation is that Ki is strongly dependent on the physicochemical properties of AAs. We hence define evolutionary AA distance of the i-th pair by Δ_U_ (i) = (U_1_-Ui)/(U_1_-U_75_) where Ui is the “universal exchangeability” given in <u>Tang et al. (2004)</u> (<u>Tang, et al. 2004</u>) (see Chen et al. (<u>Chen and Wu</u>) for an update). By this measure, Δ_U_(1) is 0 for the closest pair of AAs (Ser-Thr) and Δ_U_ (75) is 1 for the most distant pair (Asp-Tyr).

To see how well Δ_U_ may account for evolutionary rates, we separated genes into two groups: the slow group consists of the top 3 gene groups in Table 1 (0% - 60%) and the fast group consists of the last gene group (80% - 100%). *Drosophila* (*D. melanogaster* vs. *D. simulans*) and Hominoids (human vs. chimpanzee) were chosen to represent the slowest and fastest evolving taxa in our collection (Fig. 1). We found that Ki of the slow group is highly correlated with Δ_U_: the R^2^ is 0.904 in *Drosophila* and 0.706 in Hominoids (green lines in Fig. 1A-B). The R^2^ values in all other taxa are also > 0.75 (<u>Chen and Wu</u>). Hence, for 60% of the genes, there is a nearly universal relationship between Δ_U_ (i) and Ki (Fig. 1D-E). For the fastest evolving 20% of genes, the correlation decreases sharply. R^2^ among these loci is 0.798 in *Drosophila* and 0.262 in Hominoids (red lines in Fig. 1A-B).

**Figure 1.**
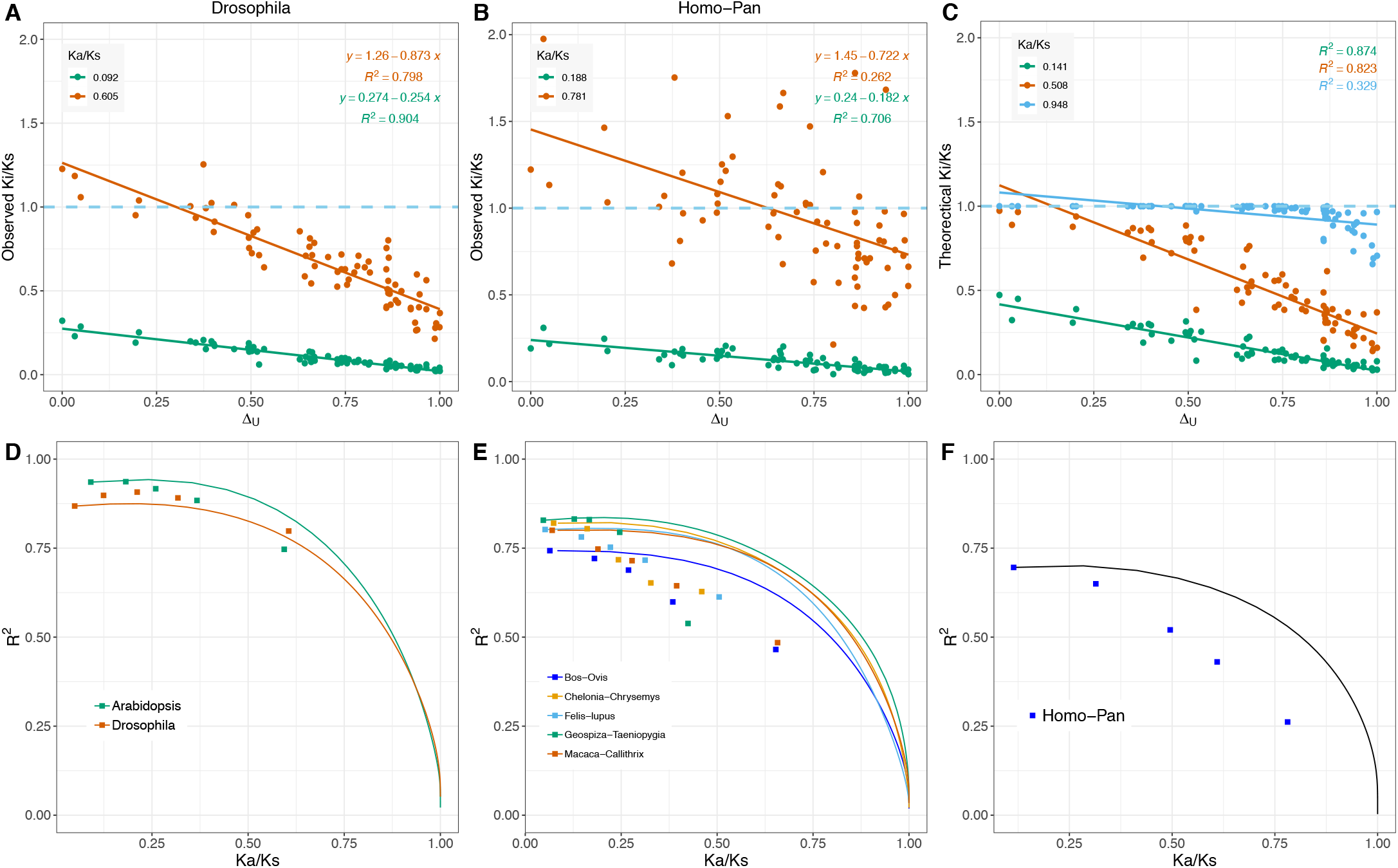
Ki/Ks vs. Δ_U_ and their correlations. (**A-B**) The relationship between Δ_U_ and Ki/Ks from rapidly and slowly evolving genes for *Drosophila* and *homo*-*pan*. Here, genes are first divided into five groups in the ascending order of their Ka/Ks ratios with equal number of non-synonymous changes in each group. Rapidly evolving genes account for the top 20% of non-synonymous changes (red points). Slowly evolving genes account for bottom 0%-60% (green points). The Ka/Ks rate ratios in the legend are the average Ka/Ks ratio for each gene group. (**C**) The theoretical changes of Ki/Ks and Δ_U_ under a model with negative selection only. As the Ka/Ks ratio increases, some AA pairs ascend to their upper limit, with Ki/Ks values nearly 1(blue points). (**D-F**) The theoretical (solid lines) and observed changes (box-shape points) of total Ka/Ks ratio and R^2^ values of their Ki against **Δ_U_**. Genes were again divided into five groups as above. The theoretical changes are based on a model with varying levels of negative selection only.

**Table 1.** The MK test for each of the five gene groups as Ka/Ks increases

| Species | Gene Group | Polymorphism |  | Divergence |  | A/S ratio |  | <sup>a</sup>F<sub>A</sub> |
| --- | --- | --- | --- | --- | --- | --- | --- | --- |
|  |  | A | S | A | S | Polym | Div |  |
| ***Drosophila***<br>(*D. melanogaster*<br>vs.<br>*D. simulans*) | 0-20% | 5,659 | 57,094 | 20,206 | 133,647 | 0.099 | 0.151 | 1.525** |
|  | 20-40% | 4,711 | 18,577 | 20,553 | 52,675 | 0.254 | 0.390 | 1.539** |
|  | 40-60% | 4,234 | 10,309 | 21,960 | 32,818 | 0.411 | 0.669 | 1.629** |
|  | 60-80% | 3,421 | 55,63 | 22,290 | 21,032 | 0.615 | 1.060 | 1.723** |
|  | 80-100% | 2,449 | 2,492 | 23,126 | 11,090 | 0.983 | 2.085 | 2.122** |
|  | ALL | 20,474 | 94,035 | 108,135 | 251,262 | 0.218 | 0.430 | 1.977** |
| **Hominoids**<br>(human vs.<br>chimpanzee) | 0-20% | 780 | 1,708 | 5,783 | 19,412 | 0.457 | 0.298 | 0.652 |
|  | 20-40% | 557 | 576 | 5,810 | 6,368 | 0.967 | 0.912 | 0.943 |
|  | 40-60% | 483 | 414 | 5,884 | 4,199 | 1.167 | 1.401 | 1.201* |
|  | 60-80% | 488 | 282 | 5,781 | 2,702 | 1.730 | 2.140 | 1.236* |
|  | 80-100% | 372 | 170 | 6,079 | 1,116 | 2.188 | 5.447 | 2.489** |
|  | ALL | 2,680 | 3,150 | 29,337 | 33,797 | 0.851 | 0.868 | 1.020 |

Polym – polymorphism; Div - divergence.
<sup>a</sup> one tailed Fisher's exact test. \* p-value < 0.05, \*\* p-value < 0.001. Here, we use Div(A/S) / Poly(A/S) for F<sub>A</sub> in lieu of Eq. (1.1) since the two expressions are nearly identical.

To further understand the underlying processes, we developed a model with variable negative selection, but without positive selection (see Methods). Fig. 1C shows simulated patterns for the relaxation of negative selection. Note that the correlation remains quite high (R2 > 0.8) as Ka/Ks exceeds 0.5. It is only when Ka/Ks approaches 1 with negative selection becoming fully relaxed, does the correlation breaks down (blue points of Fig. 1C). The observed pattern in Fig. 1A appears reasonably close to the simulated pattern in Fig. 1C, which shows the reduction in R when Ka/Ks increases. The pattern in Fig. 1B between human and chimpanzee, however, is very different from the corresponding simulations in Fig. 1C. The main reason appears to be that, in the fast group of genes, the points are often > 1 and scattered widely.

The correlations in all sampled taxa are given in Fig. 1D-F, where genes are separated into five bins by the rank order of Ka/Ks values with equal number of non-synonymous changes. The X-axis shows mean Ka/Ks ratios of each bin and the Y-axis is the R^2^ value of their Ki against Δ_U_. Simulated R^2^ values are depicted by solid lines (Fig. 1D-E; see Methods). Note that the simulations assume only negative selection in the absence of positive selection. While the simulated R^2^ agree reasonably well with the observed values in Fig. 1D (in *Drosophila* and *Arabidopsis*), the agreements in other taxa in Fig. 1E-F are much worse. Fig. 1E, which presents the five comparisons between vertebrate species (a pair of reptiles, a pair of birds, and three pairs of mammals), shows lower than expected correlations as Ka/Ks increases. Fig. 1F on the human-chimpanzee comparison shows the weakest correlations of all taxa.

Overall, the high correlation between the observed Ki and Δ_U_ starts to break down when the Ka/Ks ratio rises above 0.3 and especially above 0.5 (Fig. 1D-F). Loci with high Ka/Ks ratios are presumably less influenced by negative selection, but also possibly undergo adaptive evolution. In the next section, we attempt to separate the two effects in order to analyze the action of positive selection.

### II. Separating positive and negative selection on AA substitutions

A commonly used approach to separating positive and negative selection is the McDonald and Kreitman (MK) test (<u>McDonald and Kreitman 1991</u>). The MK test compares the Ka/Ks ratio for between-species divergence with the Pa/Ps ratio for within-species polymorphism (<u>McDonald and Kreitman 1991</u>; <u>Sawyer and Hartl 1992</u>; <u>Fay, et al. 2002</u>; <u>Smith and Eyre-Walker 2002</u>; <u>Bustamante, et al. 2005</u>; <u>Shapiro, et al. 2007</u>). The fitness advantage of nonsynonymous substitutions, F_A_, is defined as

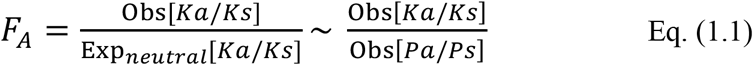

F_A_ is the observed nonsynonymous substitution relative to the expected, calibrated by Ks. Exp_*neutral*_[*Ka*/*Ks*] can then be replaced by Obs[Pa/Ps] in Eq. (1.1) if there are few advantageous variants within species. Thus, F_A_ = 1 means that nonsynonymous substitutions are not driven by positive selection. When F_A_ > 1, we interpret F_A_ as the selective advantage averaged across nonsynonymous substitutions. F_A_ < 1, which is occasionally recorded, implies weaker negative selection in the polymorphism than in the divergence data (see Discussion).

We applied the MK test to *Drosophila* and hominoids (see Fig. 1A and B). The results (Table 1) show F_A_ = 1.98 and 1.02 for the two taxa (see also Table S1, with a different frequency cutoff). When the genes are divided into five bins according to the Ka/Ks ratio, F_A_ increases steadily as Ka/Ks gets larger. Although the classification by Ka/Ks can bias the calculation of F_A_ (see Eq. 1.1), simulations show that the observed increase is faster than the expectation accounting for bias. The trend supports, albeit only indirectly, the suggestion that these fast-evolving genes are under positive selection, reducing the high correlation between Ki and Δ_U_.

We then used the MK test on each individual class of AA substitutions that differ by only one bp. As there are 75 such classes, we conducted 75 separate MK tests. For each test, Ka, Pa, and F_A_ are replaced by Ki, Pi, and Fi for i = 1, 75 (see Methods). The fitness advantage of the i-th type of AA substitution, Fi, is

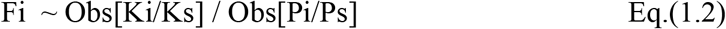

In parallel, we define the intensity of negative selection on the i-th class of AA changes as

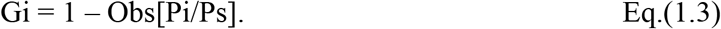

Gi is the proportion of the i-th class of codon mutations eliminated by negative selection. Because low frequency polymorphisms may contain deleterious mutations not yet eliminated (<u>Shapiro, et al. 2007</u>) and the high frequency portion of the spectrum may harbor advantageous variants (<u>Fay and Wu 2000</u>; <u>Wang, et al. 2017</u>), we followed the common practice of eliminating loci with low- and high-frequency derived alleles (see Fig. S2 and Methods for details).

With the intensity of positive and negative selection defined by Fi and Gi, we first corroborate the expectation that negative selection tolerates small-step changes. The plots of Gi against Δ_U_ in Fig. 2A-B indeed show a strong correlation with step size (see also Fig. S1 A-B). In *Drosophila*, R^2^ = 0.89. The R^2^ is lower in hominoids, but still highly significant at 0.56. The two taxa differ substantially in the strength of negative selection as can be seen on the Y-axis: Gi ranges between 0.8 and 0.99 with a mean of 0.93 in *Drosophila* but is much weaker between hominoids (as low as 0.2, with a mean of 0.68). Since the strength of negative selection determines the correlation with Δ_U_, the smaller R^2^ in hominoids is not surprising.

**Figure 2.**
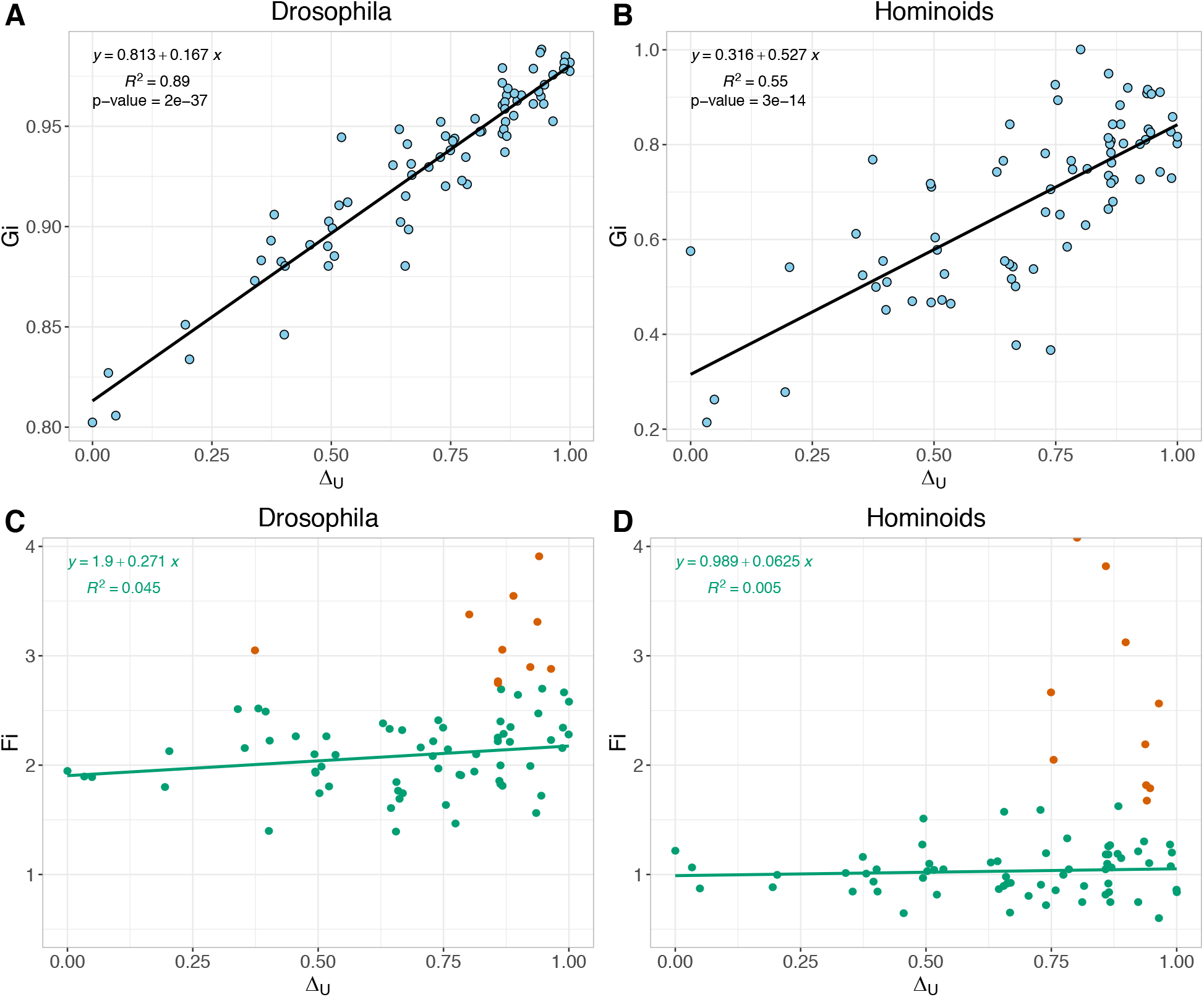
The relationship between evolutionary AA distance (Δ_U_) and selection intensity. (**A-B**) The intensity of negative selection in *Drosophila* and Hominoids. (**C-D**) positive selection intensity in *Drosophila* and Hominoids. The highest 10 Fi values are labeled by red, and the rests are green. The allele frequency cutoffs are 0.15-0.85 for *Drosophila* and 0.2-0.8 for Hominoids. In the supplementary Figure S1, different cutoffs are applied.

We now address the strength of positive selection in relation to step size, Δ_U_. In Fig. 2C-D (see also Fig. S1 C-D), the correlation can be seen as the composite of two groups of dots: Group I consists of the red dots representing the 10 highest Fi values and Group II consists of the remaining 65 green dots. The regression line for the 65 green dots is nearly flat for both *Drosophila* and hominoids. The small slopes suggest that the strength of positive selection does not depend strongly on Δ_U_. Any of the 65 AA substitutions can have a similar likelihood of being advantageous. Red dots, on the other hand, are much often favored by positiveselection and represent AA substitutions with large Δ_U_. The analysis suggests that positive selection does not favor small-step changes and the strongest adaptive signals almost always come from large-distance substitutions.

In summary, both positive and negative selection can distinguish, and act on, big-step changes better than small-step modifications. Nevertheless, there is an important difference. While negative selection appears to follow a nearly universal rule (<u>Chen and Wu</u>; <u>Tang, et al. 2004</u>), such consistency across taxa is not expected of positive selection because adaptive changes should be highly dependent on the organisms and their environments. For that reason, the similarity between Fig. 2C and D is somewhat surprising. Table 2 (and Dataset S1, see also Table S2) shows the highest 10 pairs in the Fi ranking. Five of the top 10 pairs are shared between the two taxa (p = 0.0027 with the overlap of 1.33 expected from the hypergeometric distribution). It will require extensive analyses beyond *Drosophila* and hominoids to determine if positive selection indeed favors the small subset of AAs given in Table 2.

**Table 2.**
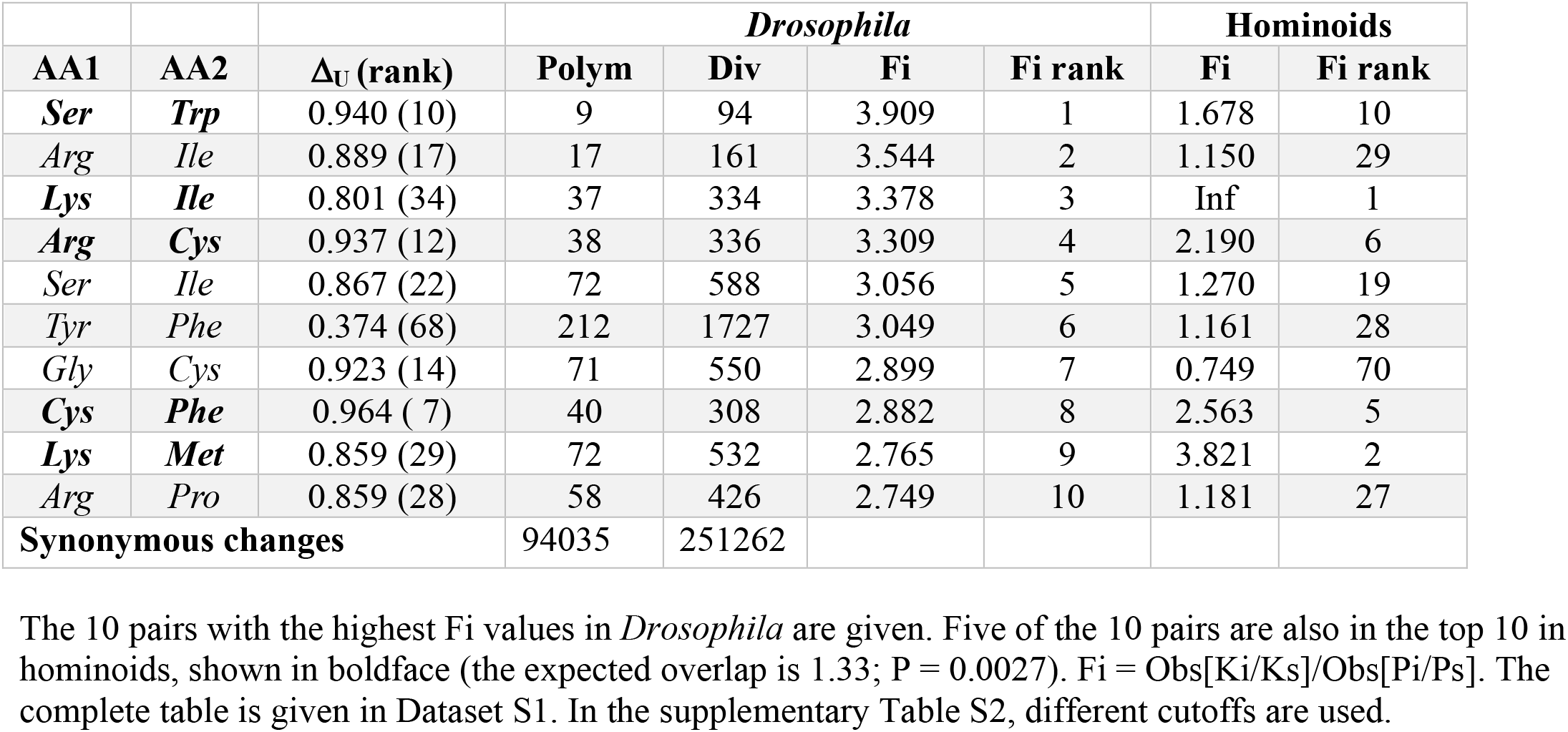
Ten AA substitutions with the highest Fi in *Drosophila* and their ranks in hominoids

| AA1 | AA2 | $\Delta_U$ (rank) | *Drosophila* | | | | Hominoids | |
| --- | --- | --- | --- | --- | --- | --- | --- | --- |
|  |  |  | Polym | Div | Fi | Fi rank | Fi | Fi rank |
| **Ser** | **Trp** | 0.940 (10) | 9 | 94 | 3.909 | 1 | 1.678 | 10 |
| **Arg** | **Ile** | 0.889 (17) | 17 | 161 | 3.544 | 2 | 1.150 | 29 |
| **Lys** | **Ile** | 0.801 (34) | 37 | 334 | 3.378 | 3 | Inf | 1 |
| **Arg** | **Cys** | 0.937 (12) | 38 | 336 | 3.309 | 4 | 2.190 | 6 |
| **Ser** | **Ile** | 0.867 (22) | 72 | 588 | 3.056 | 5 | 1.270 | 19 |
| **Tyr** | **Phe** | 0.374 (68) | 212 | 1727 | 3.049 | 6 | 1.161 | 28 |
| **Gly** | **Cys** | 0.923 (14) | 71 | 550 | 2.899 | 7 | 0.749 | 70 |
| **Cys** | **Phe** | 0.964 ( 7) | 40 | 308 | 2.882 | 8 | 2.563 | 5 |
| **Lys** | **Met** | 0.859 (29) | 72 | 532 | 2.765 | 9 | 3.821 | 2 |
| **Arg** | **Pro** | 0.859 (28) | 58 | 426 | 2.749 | 10 | 1.181 | 27 |
| **Synonymous changes** |  |  | 94035 | 251262 |  |  |  |  |

### III. Molecular evolution in small vs. large steps - A model

We have shown that the correlation between the observed Ki/Ks and Δ_U_ (i) becomes progressively weaker as the overall Ka/Ks gets higher (Fig. 1A-B). Obviously, the many Ki/Ks values above 1 require incorporation of positive selection (cf. Fig. 1B and C). An expanded model that considers the opposing dynamics of positive and negative selection is necessary. Negative selection works against large Δ_U_ (i) changes and positive selection tends to favor them. The model starts with Ka, expressed as

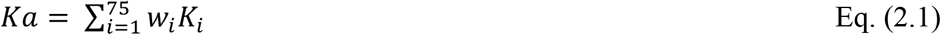

where w_i_ is the weight reflecting the number of sites available for AA exchanges of the i-th pair. The distribution of w_i_ in the *Drosophila* genome is given in Fig. 3A (top panel) where i is shown as Δ_U_ (i).

**Figure 3.**
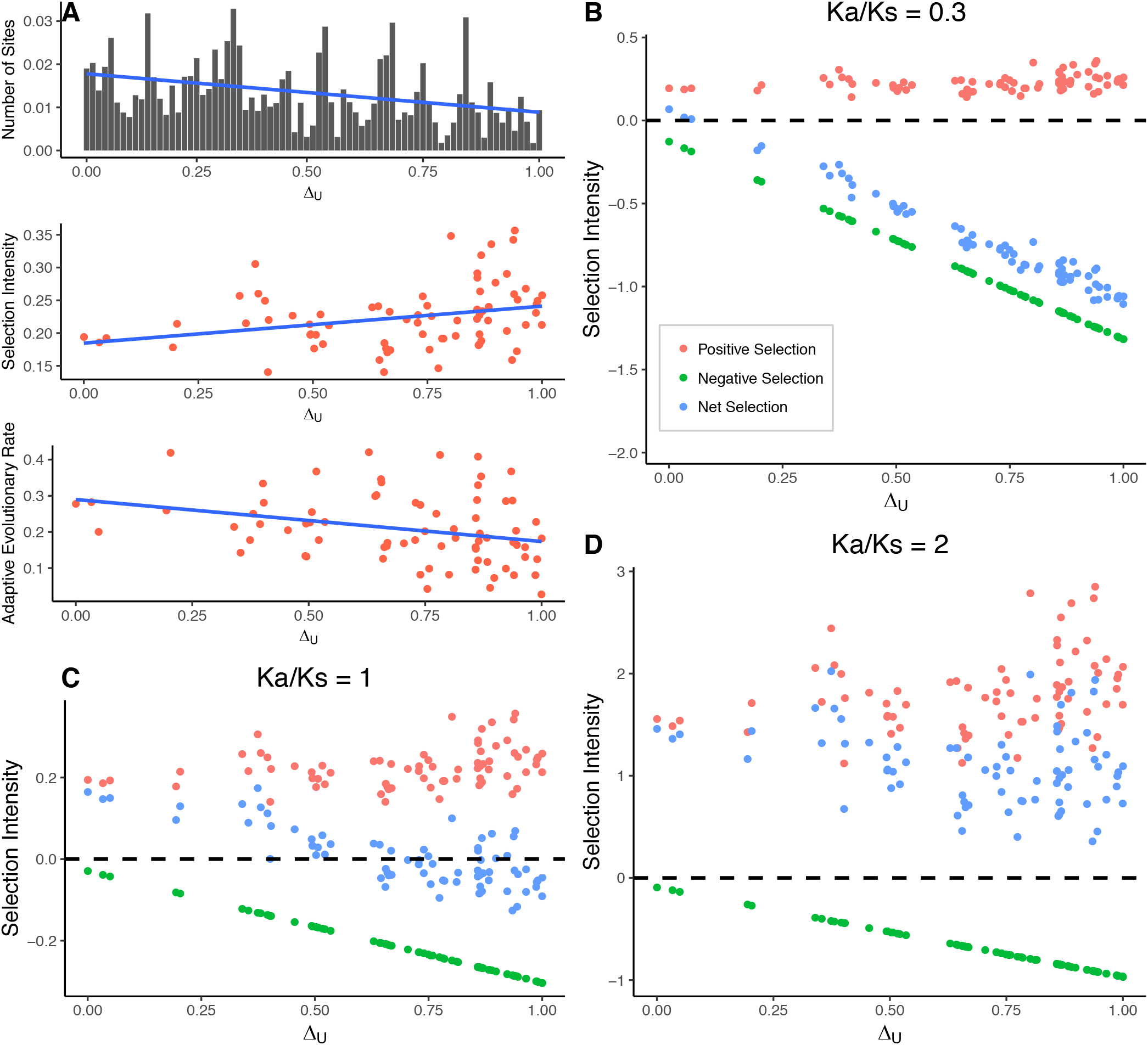
A model of selection intensity in relation to functional differences (Δ_U_). (**A**) A model with positive selection. The relative frequencies of available number of sites for each Δ_U_ (w_i_) are given in the top panel. The strength of positive selection (A_i_) relative to Δ_U_ is based on *Drosophila* data in Figure 2 (middle panel), but the adaptive evolutionary rate (the bottom panel) still depends on small-step changes due to the difference of available number of sites (w_i_) among AAs (top panel). The blue lines are the linear regression lines. (**B-D**) A model of positive selection, negative selection, and their confounding effect. There are three Ka/Ks ratios, where Ka/Ks= 0.3 (Panel **B**), Ka/Ks =1 (Panel **C**), and Ka/Ks =2 (Panel D). The dashed line denotes complete neutrality, with Ka/Ks=1. Above the dashed neutral line is the strength of positive selection (A_i_) (red points) and below the line is negative selection (B_i_, green points). The strength of negative selection (B_i_) changes linearly with functional difference (Δ_U_). The strength of positive selection (A_i_) shown in Panel A is also shared between Panel **B-C**, but the relative strength is increased in Panel **D** to achieve a higher Ka/Ks ratio. The changes of Ki/Ks relative to Δ_U_ are the result of a confounding effect between negative and positive selection (R_i_ = 1 + A_i_ − B_i_)(blue points).

We now formulate Ki (i= 1, 75) of Eq. (2.1), which is the average rate across all sites of a gene or genes.

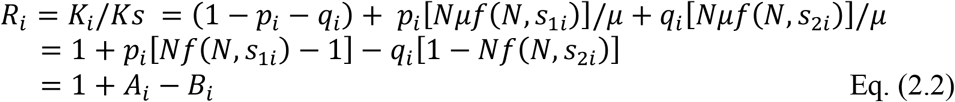

where *p_i_* and *q_i_* are the proportion of advantageous and deleterious mutations, respectively (<u>Ohta and Gillespie 1996</u>; <u>Hartl and Clark 1997</u>; <u>Li 1997</u>; <u>Chen, et al. 2018</u>). *μ* is the mutation rate and will be canceled out. In the presentation, we drop the subscript i to *s*_1_ and *s*_2_. We also use s for *s*_1_ and *s*_2_ when they are interchangeable. Then, f (N, s) = (1-e^-2s^)/(1-e^-2Ns^) is the fixation probability of a mutation with a selective coefficient s, where s can be > 0 (denoted by *s*_1_) or < 0 (*s*_2_) and N is the effective population size. *A_i_* = *p_i_*[*Nf*(*N, s*_1_) − 1] is the positive selection term and *B_i_* = *q_i_* [1 − *Nf*(*N, s*_2_)] is the negative selection term. The fixation probability of an advantageous mutation is approximately 2*s*_1_ (> 1/N) and that of a deleterious mutation is *f*(*N,s*_2_) = *ε* ~ 0. Hence, *R_i_* can be simplified as

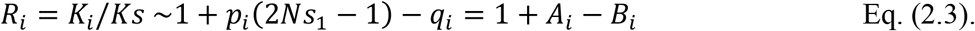

and

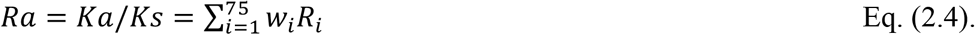

Given *p_i_*(2*Ns*_1_ − 1) > 0, it is obvious that *q_i_* ≥ 1 − *R_i_* and *p_i_* ≤ *R_i_*. Note that Ra = 0.3 is almost at the top of Ra values seen across sequenced genomes (<u>Jordan, et al. 2002</u>; <u>Drosophila 12 Genomes, et al. 2007</u>; <u>Chen, et al. 2018</u>) and, for Ra < 0.3, q > 0.7. This means that at least 70% of mutations are deleterious, allowing negative selection to dominate the overall molecular evolution.

Using Eqs. (2.3-2.4), we then quantified the relationship between *R_i_* and Δ_U_(i) for any given Ra (=Ka/Ks). In Fig. 3B-C, the Y-axis shows the strength of positive selection (*A_i_* term of Eq. 2.3; red dots above the dashed neutral line, taken from the *Drosophila* data in Fig. 2C) and negative selection (*B_i_* term; green dots below the dashed line [see the legend]). *R_i_* is the joint product, 1 + *A_i_* − *B*_i_, shown by blue dots. The X-axis shows the step size of evolution (Δ_U_).

Fig. 3B with Ra = 0.3 portrays the observed limit of a fast-evolving eukaryotic genome. Although the effects of positive and negative selection are opposite, the overall trend of *R_i_* resembles that of negative selection (blue vs. green dots). It can thus be concluded that the signature of negative selection overwhelms that of adaptive evolution in most eukaryotic genomes. For that reason, the conventional view of small-step evolution (blue dots) is hardly surprising because positive selection fails to offset the impact of negative selection. In rare cases when the strength of both selection regimes is approximately equal Ra = 1 (Fig. 3C), the overall pattern still tilts toward small-step changes. Only in the extreme case of Ra = 2 (Fig. 3D), where *A_i_* > *B_i_*, does the overall *R_i_* begin to show a weak positive correlation with step size.

### IV. Rate of adaptive evolution as a function of step size

So far we have established the efficacy of positive selection on mutations of different step sizes. The rate of adaptive evolution, however, is the product of the mutation rate and the efficacy of positive selection. Since estimation of this rate of adaptive evolution demands accounting for the influence of negative selection, it is rarely attempted (see below). The available mutational inputs of all Δ_U_ (i) in *Drosophila* are shown in the top panel of Fig. 3A, as discussed above. Large Δ_U_ changes indeed occur much less frequently than small-step mutations, a well-known property of the code table itself. In the code table, neighboring amino acids tend to be physico-chemically similar (<u>Haig and Hurst 1999</u>). The middle panel in Fig. 3A reproduces the effect of positive selection from Fig. 2C. Combining the top and middle panels of Fig. 3A, the rate of evolution in the bottom panel appears to tilt toward small-step changes. Clearly, mutational input outweighs selection efficacy in determining the overall rate. A recent study (<u>Bergman and Eyre-Walker 2018</u>) on the rate of adaptive evolution presents a pattern that is similar to (but not identical with) the bottom panel of Fig. 3A. In comparison, this study aims to show the effect of selection (middle panel), independent of the mutational input (top panel).

## Discussion

The conclusion that sets our study apart from the convention is that positive selection does not favor small-step changes (Fig. 2C-D). Although we use Δ_U_(i) as a proxy for functional changes, the analysis should have broader implications since any measure of phenotypic change can be adopted as the step size. For example, we may consider the evolved level of gene expression as the step size when studying mutations occurring in enhancers or promoters. In this sense, the small-step hypothesis for positive selection is a curious concept. When the environment changes (say, woody plants invading the intertidal zone (<u>Xu, et al. 2017</u>; <u>He, Li, et al. 2018</u>), it may require large jumps in the expression of many genes. The genome has to respond adaptively and, importantly, small-effect mutations may not be the best solution. Similarly, if the environmental changes demand modifications in the functionally important part of a protein, mutations have to fall in that part even if such alternations would have a large phenotypic effect.

In short, large-vs. small-step adaption is dictated by the environment. Small-step changes can be a good strategy for fine-tuning the phenotype, especially in an unchanging environment where the avoidance of deleterious effects is paramount. Positive selection in a new or changing environment would operate very differently. It has indeed been reported that in extreme environments or under artificial selection there is often an excess of radical amino acid substitutions (<u>Lu, et al. 2006</u>; <u>Luo, et al. 2017</u>; <u>Xu, et al. 2017</u>).

The view of small-step evolution prevails mainly because the signature of negative selection almost always dominates (see Fig. 3). Detection of the less-frequent positive selection would require filtering out the effects of negative selection. To that end, we applied the MK test to the 75 classes of AA substitutions. It should be noted that this study focuses on the *relative* Fi among the 75 classes. Hence, factors affecting theabsolute magnitude of Fi’s are of a lesser concern (<u>Fay, et al. 2002</u>; <u>Charlesworth and Eyre-Walker 2008</u>; <u>Eyre-Walker and Keightley 2009</u>; He, <u>Chen, et al. 2018</u>).

The prevalence of negative over positive selection may often have a confounding effect on the interpretation of results (<u>Wang, et al. 2017</u>). For example, previous studies have compared radical (Kr, or large-step changes) with conservative (Kc; small-step changes) AA substitutions. Kr/Kc has been shown to be higher than the overall Ka/Ks ratios (<u>Hughes, et al. 1990</u>; <u>Zhang 2000</u>; <u>Hanada, et al. 2007</u>). The trend, however, can reflect either positive selection elevating Kr or negative selection reducing Kc. In this context, Tang et al.’s two-fold rule (<u>Tang and Wu 2006</u>) is another attempt at resolving this issue. They noted that the average of the top 10 Ki classes is nearly twice the value of Ka. The accompanying study shows that the rule is correct only when Ka/Ks < 0.5 (<u>Chen and Wu</u>). This rule is hence not applicable for interpreting positive selection, which requires Ka/Ks > 1.

The concept that “nature abhors big changes” has been part of both selectionism and neutralism (<u>Fisher 1930</u>; <u>Kimura 1983</u>). This concept is only half right since it only applies to negative selection. While negative selection follows tractable common rules, the pattern of positive selection is much more taxon- and environment-dependent. The interpretation of positive selection will remain uncertain until negative selection is fully and rigorously analyzed (He, <u>Chen, et al. 2018</u>). The latter will be an immediate challenge for the study of molecular evolution.

## Materials and Methods

### The definition of Δ_U_

Functional distances between AA pairs are always based on the physicochemical properties, such as the Miyata distance (<u>Miyata, et al. 1979</u>). However, AA distances based on some physicochemical properties cannot account for all of the evolutionary exchangeability (EE) variance among amino acids. For example, AA volume differences has a prominent effect on AA properties but only accounts for 27% of the EE variance. Even the first principal component, a composite of 48 physicochemical properties, can only account for 60% of the exchangeability variance among amino acids (<u>Chen and Wu</u>).

A simple way to define a distance between AA pairs is directly based on their EE difference from genome data. These EE differences are highly conserved across a wide taxonomic range. We hence define an AA distance based on Ui, where Ui is the “universal exchangeability” given by <u>Tang et al. (2004)</u> (<u>Tang, et al. 2004</u>). Here Ui is the evolutionary exchangeability between each AA pair based on genome data. The evolutionary AA distance Δ_U_ is defined by Δ_U_ (i) =(U_1_-Ui)/(U_1_-U_75_), where U_1_ is the most exchangeable AA pair (Ser-Thr) and U_75_ is the least exchangeable (Asp-Tyr). Hence Δ_U_ (1) = 0 for the most similar pair (Ser-Thr) and Δ_U_ (75)=1 for the most dissimilar pair (Asp-Tyr). The Δ_U_ (i) is positively correlated with the functional distance.

### The definition of negative and positive selection

A widely-used method for detecting positive selection in coding regions is the MK test, which compares Ka/Ks relative to Pa/Ps (<u>McDonald and Kreitman 1991</u>). The strength of positive selection is defined as F_A_= Obs[Ka/Ks] / Obs[Pa/Ps]. F_A_ >1 indicates the existence of positive selection. In contrast, F_A_ < 1 means there is a relaxation of negative selection in polymorphism relative to divergence. Here, we do the MK test for each 75 amino acid pairs by dividing non-synonymous changes into 75 parts.

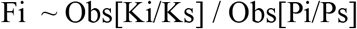

Strength of negative selection for each Δ_U_ is estimated from the polymorphism data. In the polymorphism, deleterious mutations come into the population at lower allele frequency and are then removed by negative selection. The A/S ratios of polymorphism are very high at lower allele frequencies and drop at allele prevalence increases (see Fig. S2). This drop tapers off at a certain point. We use the frequency range that has uniform Pa/Ps values in our analyses. Negative selection is then defined by

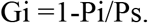

Gi is the proportion of mutations that are eliminated by negative selection.

### Multiple-alignment Data

There are eight pairs of species in Fig. 1D-E, including a pair of *Arabidopsis, Drosophila*, reptile, birds, and four pairs of mammals. Details about how to deal with the genome data can be found in our accompanying paper (<u>Chen and Wu</u>). The *Mus-Rattus* comparison was removed out as the Ui values were deduced from the comparison between *Mus-Rattus* and a pair of yeasts (<u>Tang, et al. 2004</u>).

### Polymorphism Data

Sequences of 334 *Drosophila melanogaster* lines were collected from the *Drosophila* Genome Nexus (<u>Lack, et al. 2015</u>), including 137 lines from the *Drosophila* Population Genomics Project Phase 2 (DPGP2) (<u>Pool, et al. 2012</u>) and 197 lines from DPGP3 (<u>Lack, et al. 2015</u>). After masking identical by descent and admixed regions, we assembled 1,360,772 SNPs in *Drosophila melanogaster.* Human SNPs were downloaded from the 1000 Genomes Project (<u>Consortium, et al. 2015</u>).

To estimate derived allele frequencies, adjacent species were assigned as the ancestral state. The chimpanzee and *Drosophila simulans* genomes were used as the ancestor state for human and *Drosophila melanogaster*. If the states were unknown in the adjacent species, the higher frequency alleles were assigned as ancestral states. Divergence comparisons were based on *D. melanogaster* vs. *D. similans* and human vs. chimpanzee. The divergence data of the two paired species were extracted from multiple alignment data above. There are 9,710 orthologous genes in *D. melanogaster* vs. *D. similans* comparison and and 11,571 in the human vs. chimpanzee comparison. Polymorphic mutations that were outside the orthologous genes above were removed. Especially, we masked CpG related mutations in both human polymorphism and divergence (CG => TG, CG => CA were removed).

Populations can tolerate deleterious mutations at a relatively low frequency. We removed deleterious mutations by setting a cutoff so that the A/S ratios were stable at the higher frequencies (see Fig. S2) (<u>Shapiro, et al. 2007</u>). We set two different frequency cutoff to test the influence of this choice on the results. One group is 0.2-0.8 for Hominoids and 0.15-0.85 for *Drosophila* (Results in main text). The other group is 0.2-0.95 for Hominoids and 0.15-0.95 for *Drosophila* (Results in the Supplementary Materials). Very high frequency mutations (0.95-1) were removed from further analyses because there were too many such mutations, hitting at possible positive selection in this class (<u>Fay and Wu 2000</u>; <u>Wang, et al. 2017</u>).

### Relaxation of negative selection models

Fitness effects of new deleterious mutations were drawn from the exponential distribution. Most deleterious mutations are under strong negative selection, thus the fitness is nearly 0. A small fraction of mutations are under weak negative selection with fitness far greater than 0.

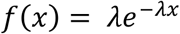

Here, f(x) is the frequency of mutations. The x is the fitness of new deleterious mutations. The fixation probability of such mutations is the cumulative fitness from the least fit to current fitness, thus

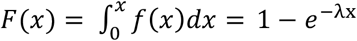

Ka/Ks is the fixation probability of new non-neutral relative to neutral mutations. Thus F(x) = Ka/Ks in the case where only deleterious mutations are under consideration.

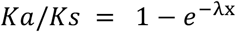

Here, we connect the relationship between the fitness of a mutation x and the Ka/Ks ratio. As the fitness of a new mutation is in the inverse relationship with the strength of negative selection, mutations under strong negative selection undergo significant fitness reduction. The equation above thus links the strength of negative selection and the Ka/Ks ratio. When mutations are under strong negative selection, x ~0 and Ka/Ks is nearly 0. When constraint is absent, x ~ *infinite* and Ka/Ks is 1.

Thus we define the fitness of new deleterious mutations x and Ki/Ks for different Δ_U_(i=1:75) as follows:

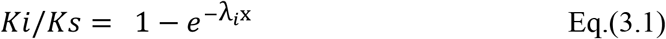

We simulated relaxation of negative selection based on Eq. (3.1). First, we set x as a constant value and obtained *λ_i_* (i=1:75) for each Δ_U_, under the lowest total Ka/Ks ratio situation from the real data. We get the Ki/Ks values for each Δ_U_ of the slowest evolving genes accounting for 20% of all non-synonymous changes and calculate the corresponding *λ_i_* (i=1:75) from Eq. (3.1). Relaxation of negative selection was simulated by predicting Ki/Ks changes as the total Ka/Ks ratio increased. We simply increased the fitness x and calculated the corresponding theoretical Ki/Ks values (i= 1:75) based on Eq. (3.1). By changing the magnitude of x, the theoretical Ki/Ks values under every possible total Ka/Ks ratio were calculated. In the complete absence of constraint, λix values were nearly *infinite* and theoretical Ki/Ks reached their upper limit, Ki/Ks ~1 for most Δ_U_.

## Acknowledgements

We would like to thank Pei Lin and members of Wu Lab for discussions and advices. This work was supported by the National Natural Science Foundation of China (91731301 and 31730046), the 985 Project (33000-18831107), and National Key Basic Research Program of China (2014CB542006).

