## Supplementary for "Molecular evolution in large steps - Codon substitutions under positive selection"

**Table S1 – The MK test for each of the five gene groups with increasing Ka/Ks**

| Species | Gene Group | Polymorphism | | Divergence | | A/S ratio | | $^{a}F_A$ |
|---|---|---|---|---|---|---|---|---|
| | | A | S | A | S | Polym | Div | |
| **Drosophila** **(D. mel vs. D. simulans)** | 0-20% | 6,153 | 63,524 | 20,206 | 133,647 | 0.097 | 0.151 | 1.561** |
| | 20-40% | 5,189 | 20,747 | 20,553 | 52,675 | 0.250 | 0.390 | 1.560** |
| | 40-60% | 4,721 | 11,481 | 21,960 | 32,818 | 0.411 | 0.669 | 1.627** |
| | 60-80% | 3,818 | 6,242 | 22,290 | 21,032 | 0.612 | 1.060 | 1.733** |
| | 80-100% | 2,837 | 2,789 | 23,126 | 11,090 | 1.017 | 2.085 | 2.050** |
| | ALL | 22,718 | 104,783 | 108,135 | 251,262 | 0.217 | 0.430 | 1.985** |
| **Hominoids** **(human vs. chimpanzee)** | 0-20% | 884 | 2,022 | 5,783 | 19,412 | 0.437 | 0.298 | 0.681 |
| | 20-40% | 648 | 697 | 5,810 | 6,368 | 0.930 | 0.912 | 0.981 |
| | 40-60% | 580 | 466 | 5,884 | 4,199 | 1.245 | 1.401 | 1.126* |
| | 60-80% | 576 | 334 | 5,781 | 2,702 | 1.725 | 2.140 | 1.241* |
| | 80-100% | 437 | 191 | 6,079 | 1,116 | 2.288 | 5.447 | 2.381** |
| | ALL | 3,125 | 3,710 | 29,337 | 33,797 | 0.842 | 0.868 | 1.031 |

Like Table 1, different allele frequency cutoffs are applied, with 0.15-0.95 for *Drosophila* and 0.2-0.95 for Hominoids.
Polym – polymorphism; Div - divergence.
[a] one tailed Fisher's exact test. * p-value < 0.05, ** p-value < 0.001. Here, we use Div(A/S) / Poly(A/S) for $F_A$ in lieu of Eq. (1.1) since the two expressions are nearly identical.

**Table S2 – Ten AA substitutions with the highest Fi in *Drosophila* and their ranks in hominoids**

| | | | Drosophila | | | | Hominoids | |
|---|---|---|---|---|---|---|---|---|
| AA1 | AA2 | $\Delta_U$ (rank) | Polym | Div | FI | FI rank | FI | FI rank |
| ***Ser*** | ***Trp*** | 0.940 (10) | 11 | 94 | 3.564 | 1 | 1.976 | 8 |
| ***Lys*** | ***Ile*** | 0.801 (34) | 40 | 334 | 3.482 | 2 | 3.403 | 1 |
| *Arg* | *Cys* | 0.937 (12) | 41 | 336 | 3.418 | 3 | 1.587 | 12 |
| *Arg* | *Ile* | 0.889 (17) | 20 | 161 | 3.357 | 4 | 1.354 | 16 |
| *Ser* | *Ile* | 0.867 (22) | 77 | 588 | 3.185 | 5 | 1.197 | 23 |
| *Tyr* | *Phe* | 0.374 (68) | 236 | 1727 | 3.052 | 6 | 1.003 | 48 |
| *Arg* | *Pro* | 0.859 (28) | 61 | 426 | 2.912 | 7 | 1.270 | 18 |
| *Gly* | *Cys* | 0.923 (14) | 79 | 550 | 2.903 | 8 | 0.849 | 65 |
| ***Lys*** | ***Met*** | 0.859 (29) | 78 | 532 | 2.844 | 9 | 3.000 | 3 |
| ***Arg*** | ***Met*** | 0.898 (16) | 33 | 219 | 2.768 | 10 | 2.452 | 4 |
| | | Synonymous | 104783 | 251262 | | | | |

The 10 pairs with the highest Fi values in *Drosophila* are given. Four of the 10 pairs are also in the top 10 in hominoids, shown in boldface (the expected overlap is 1.33; P = 0.024). Like Table 2, different allele frequency cutoffs are applied, with 0.15-0.95 for *Drosophila* and 0.2-0.95 for Hominoids.

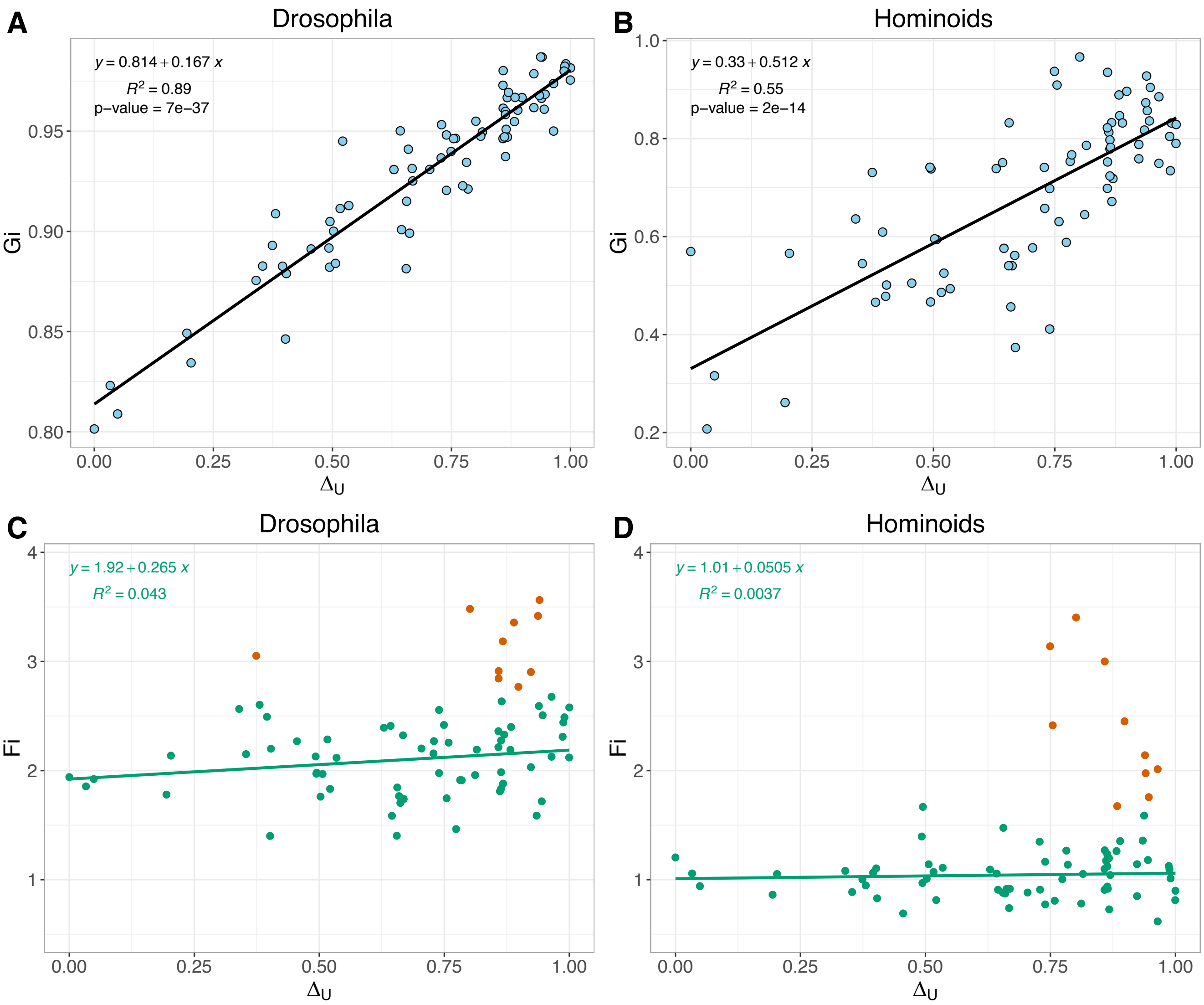


**Figure S1 | Relationship between evolutionary AA distance ($\Delta_U$) and selection.** (**A-B**) Negative selection intensity in *Drosophila* and Hominoids. (**C-D**) positive selection intensity in *Drosophila* and Hominoids. The highest 10 Fi values are labeled in red, the rest are green. Like Figure 2, different allele frequency cutoffs are applied, with 0.15-0.95 for *Drosophila* and 0.2-0.95 for Hominoids.

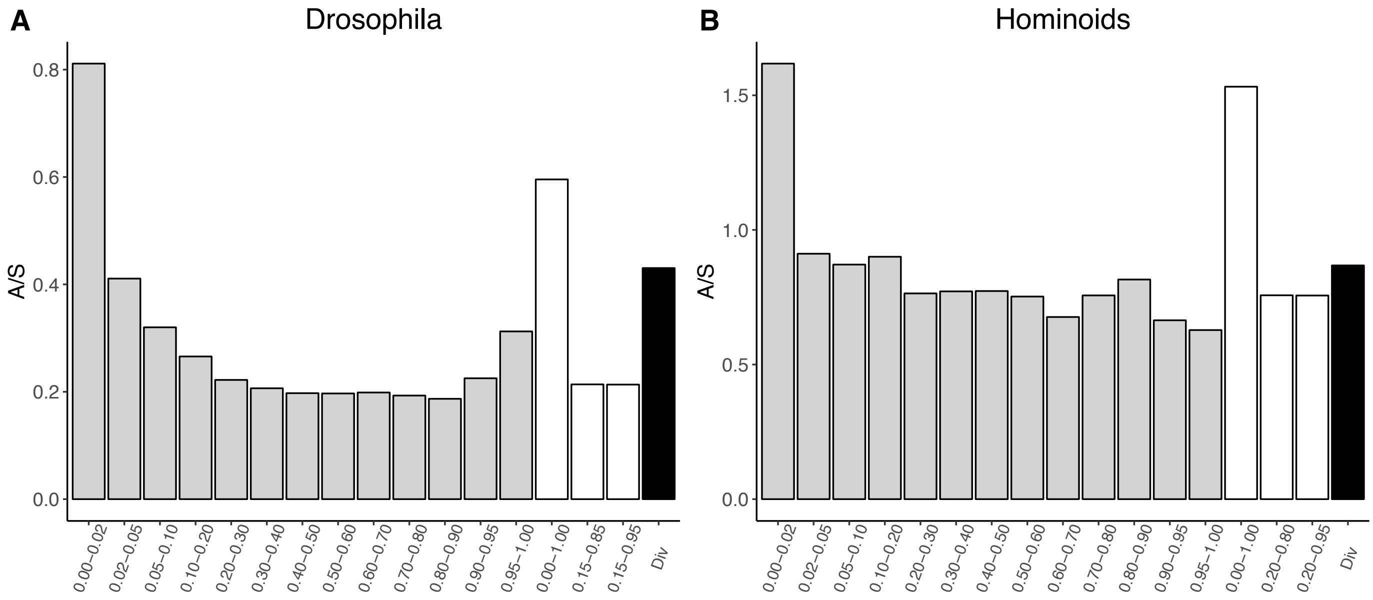


**Figure S2| A/S ratio of polymorphism as a function of derived allele frequency.** The gray bars are the A/S ratio from 0-1, separated by their derived allele frequency. The open bars are A/S for all polymorphism data and two frequency cutoffs used in the paper. The black bars are the divergence A/S ratio between *D. melanogaster* vs. *D. simulans* (**A**) and human vs. chimpanzee (**B**).
